# Immunodominance is a poor predictor of vaccine-induced T follicular helper cell quality

**DOI:** 10.1101/2025.07.21.666029

**Authors:** HX Tan, MZM Zheng, K Wragg, L Murdiyarso, D Pilapitiya, A Kelly, R Esterbauer, C Gonelli, AK Wheatley, JA Juno

## Abstract

Rational engineering of vaccine immunogens to focus B cell responses on potently neutralizing epitopes is a promising approach to improve the potency, breadth and durability of viral vaccines. Such strategies, however, can compromise vaccine immunogenicity through the unintended exclusion of CD4+ T cell epitopes, which are critical for the development of T follicular helper (TFH) cells and to support high affinity antibody production. Using a prototypic influenza HA stem immunogen lacking effective CD4+ T cell help in BL6 mice, we interrogated the minimal requirements for T cell help needed to drive serological responses to vaccination. We find that priming of naïve CD4 T cells is markedly efficient, however the immunodominance of a given CD4 T cell epitope is not predictive of the propensity to provide high quality help to antigen-specific B cells. In the context of soluble antigens, provision of a single MHC class II epitope is sufficient to drive robust germinal centre responses and serum IgG titres. However not all CD4 epitopes provide equivalent levels of B cell help, despite priming comparable numbers of antigen-specific CD4 T cells. Finally, we show multimerizing and arraying antigens on nanoparticle scaffolds unlocks highly subdominant, near-undetectable CD4 T cell helper responses to support a T-dependent antibody response. Our findings emphasize the importance of CD4+ T cell help for programing robust and durable humoral immunity, and provide crucial insights to guide the rational incorporation of favorable T cell epitopes into vaccines.

## Introduction

The development of effective vaccines has fundamentally reshaped the ability of human populations to contain and control infectious diseases. While historically vaccines were derived from attenuated or killed whole pathogens, more recent vaccine development efforts have focused upon identification of the most protective “subunits” for inclusion as vaccine immunogens. This was extended during the COVID-19 pandemic, where rational vaccine design was used to engineer the SARS-CoV-2 spike immunogens in protein, mRNA or viral-based vaccines to structurally stabilize^1,2^ and/or prevent cleavage^3,4^ of the spike trimer. Analogous modifications of the F protein are incorporated into recently approved vaccines for RSV^5^.

Further efforts to concentrate immunity upon conserved or potently neutralizing domains have seen re-engineering of vaccine immunogens to the level of isolated protein subdomains such as the SARS-CoV-2 RBD^6,7^ or stem region of influenza HA^8^. While such small protein targets effectively focus immune recognition by B cells, increasing evidence suggests this might come at a cost to immunogenicity, seen in both pre-clinical models^9,10^ and human clinical trials^11,12^.

Outside of the stochasticity of vaccine delivery, the immunogenicity of vaccine antigens varies, likely reflecting a combination of protein intrinsic factors and host genetics. We and others have identified that a paucity of CD4 helper T cell epitopes is one important factor constraining vaccine immunogenicity^9,10,13,14^. Unlike B cells, whose immunoglobulin receptors scan and engage the near infinite diversity of conformational epitopes on protein surfaces, CD4+ T cells can only recognize linear peptide epitopes in the context of a host MHC-II. This heightened restriction limits the absolute number of T cell epitopes within any given immunogen, with distribution being uneven and individualistic^15^. Reducing the size of a vaccine immunogen therefore has the potential to concomitantly shrink the pool of available CD4+ T cell epitopes, leaving some antigens poorly recognized even in diverse human populations^16-18^. In extreme examples, exemplified by model HA stem^9^ or HEL^19^ immunogens in genetically inbred C57BL/6 mice, a lack of CD4 T cell epitopes can render immunization immunologically silent.

Various strategies have been reported to augment CD4 T cell priming, T follicular helper cell (TFH) differentiation, and subsequent vaccine immunogenicity, but generalized applicability, comparative utility, and mechanisms of action remain unclear. Presentation of vaccine antigens on nanoparticle scaffolds enhances BCR recognition and signaling^20^, but can also augment CD4 T cell help via epitopes localized within the scaffold^21^. Similarly, covalent coupling of poorly immunogenic proteins to carrier proteins or alternate sources of T cell help can supplement the available antigen-specific CD4 T cell pool^9,22^, analogous to the carrier proteins essential for driving antibody responses to pneumococcal carbohydrates in childhood vaccines^23^. However the impacts of epitope specificity or relative immunodominance of polyclonal CD4+ T cell populations upon the capacity to provide help to germinal centre (GC) B cells is poorly understood.

Here, we made use of the HA stem-C57BL/6 model^9^ to address the features of CD4 T cell immunity that drive potent humoral immune responses to vaccination. We find evidence for a spectrum of epitope-level regulation of GC initiation, with some CD4 T cell specificities unable to support antigen-specific B cell proliferation and IgG production. Critically, epitope immunodominance did not necessarily predict the quality of help provided by antigen-specific CD4 T cells: some immunodominant epitopes exhibited no helper capability, while subdominant responses provided effective help, with even CD4 T cells at near-undetectable levels successfully contributing to GC formation in the context of multimerized antigen. Overall, these data demonstrate that while vaccine immunogenicity is critically dependent on CD4 T cell availability, the capacity to support humoral responses is not universal to all CD4 T cell epitopes, with implications for the rational design of small, engineered vaccine immunogens.

## Results

### Multimeric display of HA stem antigens on self-assembling ferritin nanoparticles compensates for highly subdominant CD4 help

We previously established primary vaccination of BL/6 mice with a soluble HA stem protein fails to elicit an antigen-specific GC B cell response or production of serum IgG due to a lack of available CD4 T cell epitopes^9^. Presentation of stem on self-assembling ferritin nanoparticles enhances its immunogenicity^24,25^ but the source of T cell help in this system is unclear, as BL/6 mice are reported to lack ferritin-specific CD4 T cells^21^. We find that stem-ferritin nanoparticles (Stem-Fe) are robustly immunogenic, eliciting significantly higher stem IgG titres than soluble stem trimers (p=0.0001; Fig1A) and greater numbers of bulk and stem-specific GC B cells (p=0.0001 and p=0.02, respectively; Fig 1B-C, gating in Supplementary Fig 1). Using confocal imaging and flow cytometry, we confirmed the GL7^hi^CD38^lo^ B cell populations in stem-Fe vaccinated mice reflected *bona fide* GC structures. GL7 expression was correctly localized to GC-like structures within the follicle (Supplementary Fig 2A), and canonical light zone and dark zone GC B cell populations were observed in both full-length HA (HA-FL) and stem-Fe animals (Supplementary Fig 2B), indicative of a T-dependent GC reaction.

**Figure 1.**
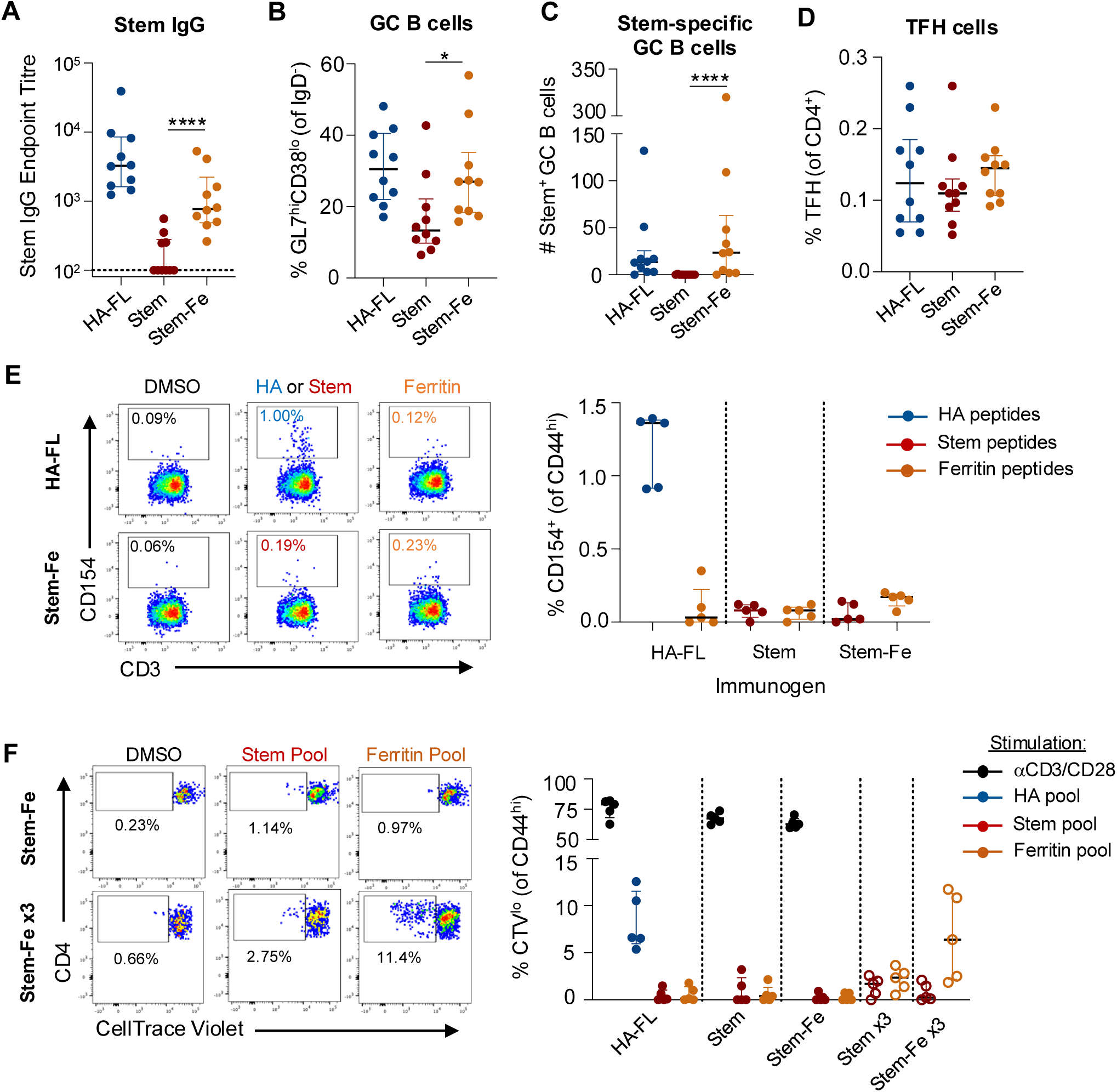
Nanoparticle display enhances stem immunogenicity in the absence of prominent CD4 T cell help. Mice were vaccinated with 5mg of full-length HA protein (HA-FL), HA-stem protein, or stem-ferritin nanoparticles (Stem-Fe) co-formulated 1:1 with Addavax adjuvant. Serum and draining LN were collected at day 14 post-vaccination. **(A)** Serum endpoint titres of stem-specific IgG (N=10/group). **(B)** Frequency of GL7^hi^CD38^lo^ GC B cells, **(C)** number of stem-specific GC B cells, or **(D)** frequency of CXCR5^hi^PD-1^hi^ TFH in the draining LN (N=10/group). **(E)** Representative staining and frequency of CD154^+^ stem or HA-specific memory CD4 T cells following *in vitro* stimulation. Frequencies are background subtracted based on the DMSO control (N=5/group). **(F)** CD4^+^ T cell proliferation following *in vitro* peptide stimulation with HA, stem or ferritin peptide pools. Splenocytes were harvested from vaccinated mice at day 14, or at day 79 after 3 vaccinations (N=5/group). Lines indicate median and IQR. Statistics assessed by Mann-Whitney test comparing Stem and Stem-Fe groups. *p<0.05, ****p=0.0001

Given the lack of identifiable CD4 T cell epitopes within the stem antigen and the limited impact of stem-Fe vaccination on total TFH frequency (Fig 1D), we assessed the magnitude of ferritin-specific responses. We observed limited CD4 T cell responsiveness upon *in vitro* re-stimulation with either stem or ferritin peptide pools (Fig 1E). Similarly, use of a proliferation assay for sensitive detection of any low-frequency populations of stem- or ferritin-specific T cells revealed only sporadic, low-level proliferation when compared to the vehicle control (Fig 1F). To further amplify antigen-specific T cell frequencies, we primed and then boosted mice twice on days 21 and 42. Three out of five Stem-Fe vaccinated animals exhibited clear proliferative responses to the ferritin peptide pool (Fig 2F), suggesting highly subdominant epitopes within ferritin could be the source of T cell help supporting Stem-Fe immunogenicity.

**Figure 2.**
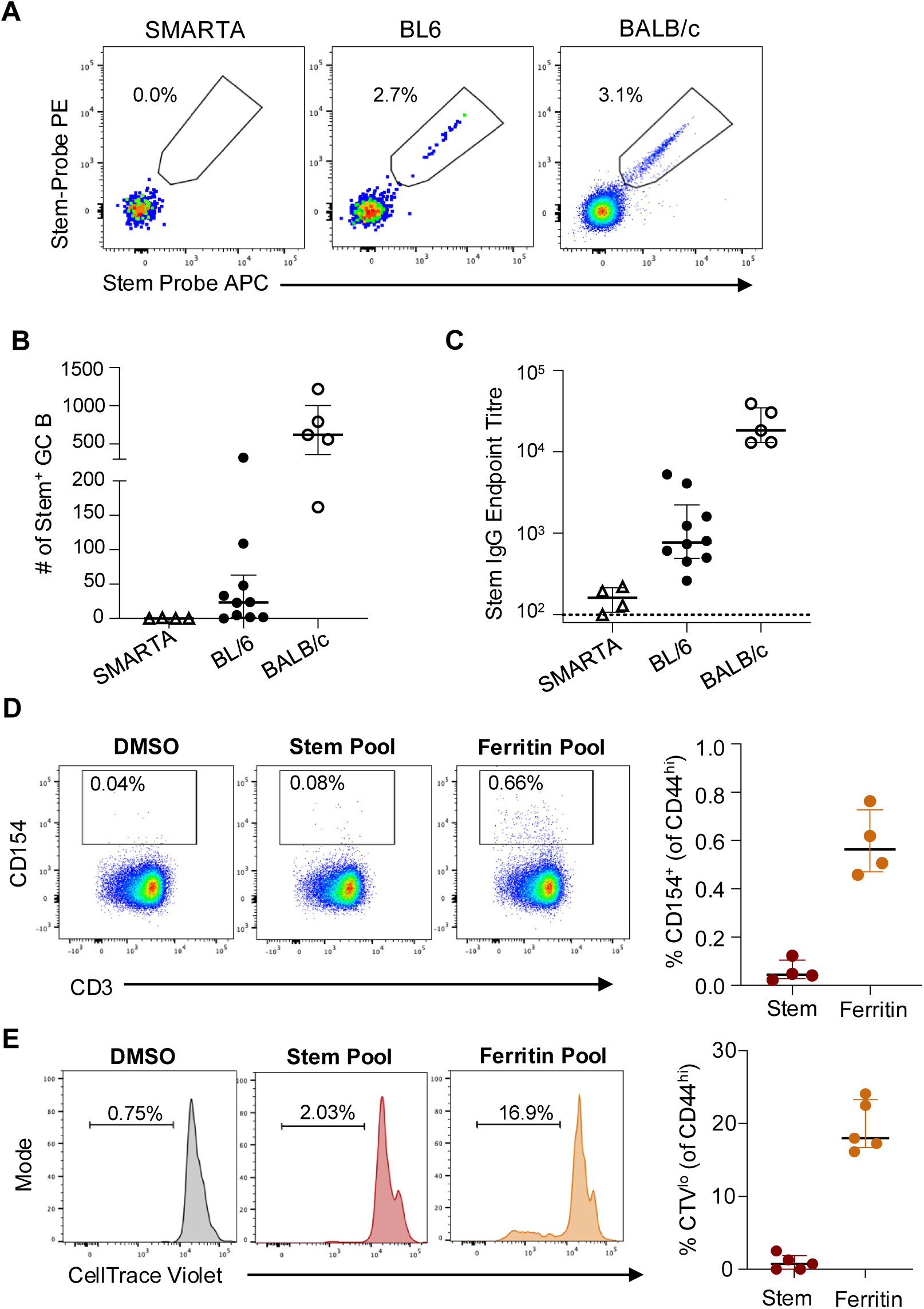
Stem-Fe nanoparticle immunogenicity in SMARTA and BALB/c mice. Stem-specific **(A)** GC B cell frequencies, **(B)** GC B cell counts or **(C)** IgG titres at day 14 following Stem-Fe vaccination of SMARTA (N=4), WT BL6 (N=10) or BALB/c (N=5) mice. **(D)** Stem- or ferritin-specific CD4^+^ T cell responses in the draining LN of Stem-Fe vaccinated BALB/c mice at day 14 (n=4). **(E)** CD4^+^ T cell proliferation in BALB/c mice following *in vitro* peptide stimulation with DMSO, stem or ferritin peptide pools. Splenocytes were harvested from vaccinated mice at day 14 (N=5/group). Lines indicate median and IQR.

If ferritin-specific CD4 help below the detection limit of conventional T cell assays was responsible for supporting a primary GC reaction, we reasoned that the Stem-Fe antigen should not be immunogenic in transgenic (tg) BL/6 mice that express a single TCR. Conversely, animals with high frequencies of ferritin-specific T cells (i.e. BALB/c mice^21^) should produce superior serological responses. Stem-Fe vaccination of SMARTA tg (BL/6 background), wild-type (WT) BL/6 and WT BALB/c animals demonstrated a striking immunogenicity gradient, with negligible numbers of stem-specific GC B cells (Fig 2A-B) or IgG (Fig 2C) detected in SMARTA mice, while responses in WT BALB/c were more than 20-fold higher compared to WT BL/6 (Fig 2A-C). We confirmed that ferritin-specific CD4 T cells were readily detectible in BALB/c mice by both standard restimulation (Fig 2D) and proliferation assays (Fig 2E), demonstrating that the ferritin core provides substantially more CD4 T cell help in BALB/c versus BL/6 mice.

Overall, when stem is arrayed on the surface of a nanoparticle, the low level of ferritin-specific CD4 T cell help available in BL/6 mice becomes sufficient to support a stem-specific GC response that is otherwise absent following soluble stem vaccination.

*The TFH repertoire of full-length HA is limited in breadth and dominated by a single epitope* The gradient of Stem-Fe immunogenicity across SMARTA, BL/6 and BALB/c mice strains suggests that availability of CD4 T cell help acts as a rheostat that directly tunes the magnitude of the GC and resultant serological response. While we have established the near-complete lack of CD4 help in stem, the breadth and specificity of epitopes that successfully support the immunogenicity of the full-length HA (HA-FL) protein requires detailed mapping.

To maximize the number of TFH for screening, BL/6 mice were infected with a sublethal dose of PR8 influenza virus and mediastinal LN (mLN) which drain the site of infection were collected on day 14. LN cell suspensions were stimulated *in vitro* with a matrix of 20 peptide pools spanning the HA protein (Supplementary Fig 3A), with candidate immunogenic peptides selected by identifying pools that elicited a CD154 response greater than the DMSO control (Supplementary Fig 3B). Individual peptide screening identified 14 potential hits, which we subsequently tested for consistent recognition across multiple animals. Nine peptides elicited CD4 T cell responses in at least 3 of 5 mice, with 4 putative epitopes identified for the CXCR5^+^PD-1^+^ CD4 T cell population (enriched for pre-TFH/TFH cells; Fig 3A). HA_91-107_ was identified as highly immunodominant, with a median of 2.6% of TFH specific for this single peptide. Three other epitopes elicited lower magnitude responses: HA_301-323_ (two overlapping peptides spanning HA_301-317_ and HA_307-323_), HA_115-131_, and HA_523-539_ (Fig 3A). A similar hierarchy was observed within the total CD4+ T cell population (Supplementary Fig 3C), with no immunogenic peptides located within the HA-stem domain as expected.

**Figure 3.**
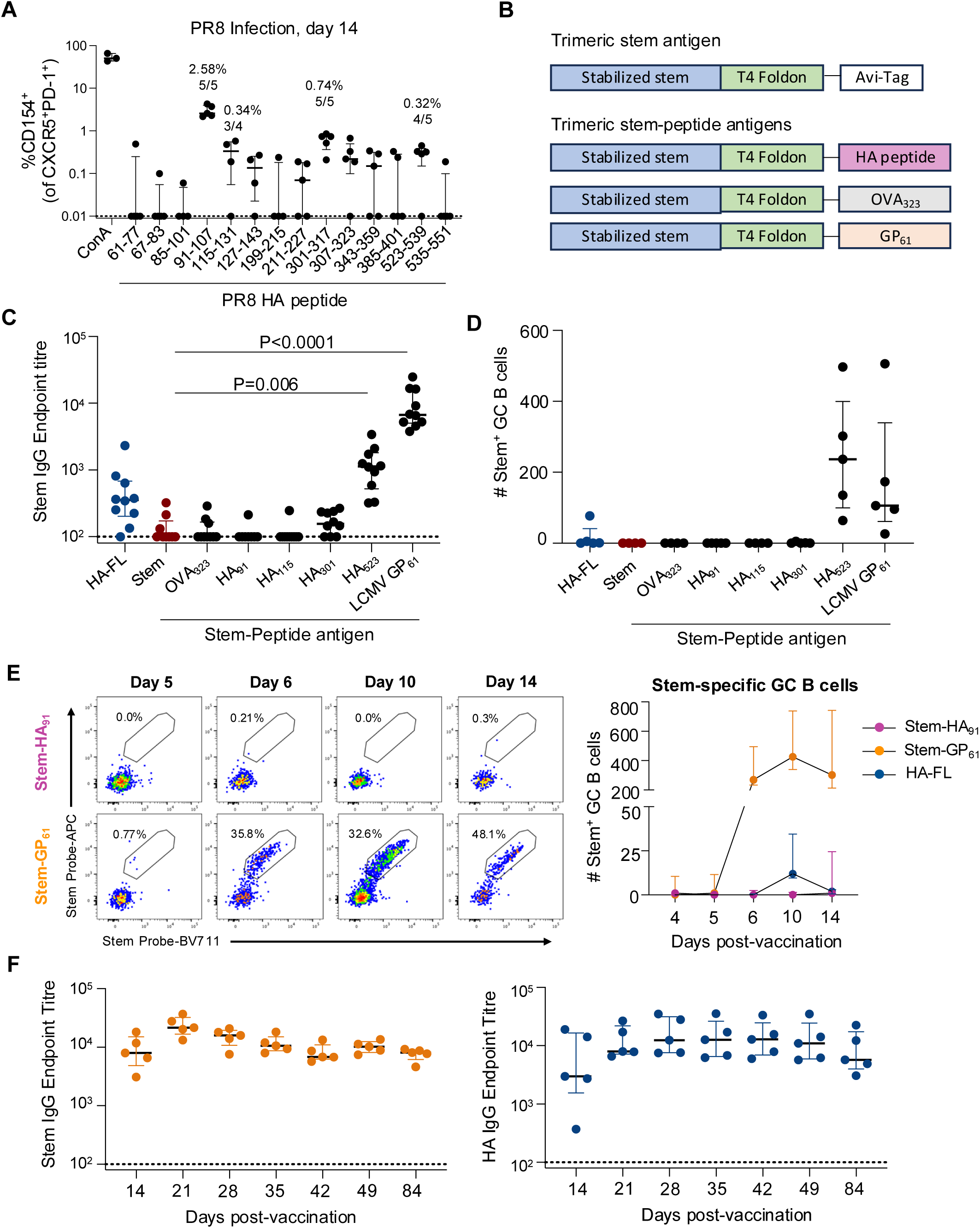
Rescue of stem immunogenicity by genetically fused CD4 T cell epitopes. **(A)** Identification of immunogenic peptides following intranasal infection with PR8 virus. Mediastinal LN were collected on day 14 and restimulated in vitro with DMSO, ConA or indicated peptides (N=4-5 mice per peptide). Immunogenic peptides are labelled with the median response and number of responding mice. **(B)** Design of trimeric stem antigens covalently linked to HA-derived peptides or prototypic OVA_323_ and GP_61_ peptides. **(C)** Stem-specific IgG endpoint titres at day 14 post-vaccination with 5ug of HA-FL, stem, or stem-peptide antigens formulated with Addavax (N=7-10 per group). Statistics assessed by Kruskal-Wallis test with Dunn’s post-test compared to the stem control. **(D)** Number of stem-specific GC B cells (GL7^+^CD38^lo^) at day 14 post-vaccination (N=4-5 per group). **(E)** Longitudinal tracking of stem-specific GC B cells in the draining LN at days 4, 5, 6, 10 and 14 post-vaccination (N=5 per group). **(F)** Durability of antigen-specific serum IgG following stem-GP_61_ or HA-FL vaccination (N=5 per group). Symbols or lines indicate median with IQR.

To confirm whether these same peptides were immunogenic in the context of vaccination, we intramuscularly immunized mice with 5μg of soluble HA-FL protein in Addavax adjuvant. HA_91_ epitope was again immunodominant, accounting for 74% of the CD154 response to the full HA peptide pool (median of 7.85% CD154^+^ for HA pool versus 5.76% for HA_91_ peptide) (Supplementary Fig 3D).

### Addition of an MHC class II epitope to soluble stem protein rescues stem-specific IgG

Given the wide spectrum of TFH responses capable of modulating serological outcomes, from low-level ferritin-specific help for nanoparticle-driven GCs to the heavy dominance of HA_91_ in the native HA-specific T cell pool, we further sought to clarify the intrinsic “helpfulness” of discrete CD4+ T cell epitopes. We tested whether restoration of a single epitope onto the stem vaccine antigen could prime cognate CD4+ T cells in vivo and thereby improve the stem-specific IgG response. Variants of the trimeric stem immunogen were developed with incorporation of a single peptide epitope sequence adjoined at the C-terminus of the T4 foldon domain (Fig 3B). We produced immunogens incorporating each peptide mapped from HA (HA_91_, HA_115_, HA_301_ and HA_523_) as well as prototypic IA^b^-restricted peptides OVA_323_ (‘OTII’) and LCMV GP_61_ (‘SMARTA’).

WT BL/6 mice were vaccinated with a single 5μg dose of HA-FL, stem, or stem variant antigens formulated in Addavax. On day 14, stem IgG titres were assessed by ELISA and stem-specific GC B cells quantified in the dLN. Only two antigens, stem-HA_523_ and stem-GP_61_, were capable of eliciting stem-specific IgG and GC B cells at levels above the stem control (p=0.006 and p<0.0001, respectively; Fig 3C-D). The failure of the immunodominant HA_91_ epitope to rescue the stem response prompted us to examine the biogenesis of the GC response from days 4-14 post-vaccination. Stem-HA_91_ vaccinated animals failed to form a stem-specific GC B cell population at any timepoint, while stem-GP_61_-immunised animals exhibited a robust stem-specific GC response evident from day 6 onwards (Fig 3E). At peak, antigen-specific GC B cell numbers were 33-fold higher in Stem-GP_61_ vaccinated animals compared to HA-FL, with the serological response demonstrating similar durability for both antigens (Fig 3F).

### HA_91_-specific TFH cell fail to populate the germinal centre

Despite HA_91_ dominance in the native CD4+ T cell response to HA-FL vaccination and PR8 infection, incorporation of the HA_91_ peptide failed to rescue stem immunogenicity. Possible mechanisms could include a peptide processing defect in the engineered stem immunogen that prevented or reduced naïve CD4 T cell priming by DCs, or alternatively a qualitative defect in TFH differentiation and/or T cell:B cell interactions. To address this, we produced an IA^b^/HA_91_ tetramer that facilitated tracking of antigen-specific T cells. In both PR8 infection and HA vaccination, staining of the draining LN with tetramer confirmed the expansion of a prominent CD44^hi^ epitope-specific CD4 T cell population (Supplementary Fig 4A). A similar tetramer successfully identified GP_61_-specific T cells in dLN following LCMV GP immunization (Supplementary Fig 4A).

Using the HA_91_ and GP_61_ tetramers, we compared epitope-specific CD4 T cell numbers and phenotypes at days 4, 5, 6, 10 or 14 post-vaccination with either stem-HA_91_ or stem-GP_61_ (gating in Supplementary Fig 4B). Both antigens readily primed CD4 T cells, evidenced by the expansion of a tetramer^+^ CD44^hi^ population (Fig 4A) with progressive maturation from a CD62L^hi^ to CD62L^lo^ phenotype (Supplementary Fig 4C). Total numbers of HA_91_- or GP_61_-specific T cells in the dLN were comparable, although the stem-HA_91_ antigen primed significantly more tetramer^+^ cells at day 5 compared to HA-FL protein (Fig 4A). Nevertheless, these populations followed different TFH differentiation trajectories over the subsequent 11 days. The small numbers of CXCR5^hi^PD-1^hi^ TFH-like GP_61_-specific cells seen on day 4 became a substantial TFH population by day 6, when median numbers of antigen-specific TFH were 4.2-fold greater for GP_61_ compared to HA_91_ (Fig 4B).

**Figure 4.**
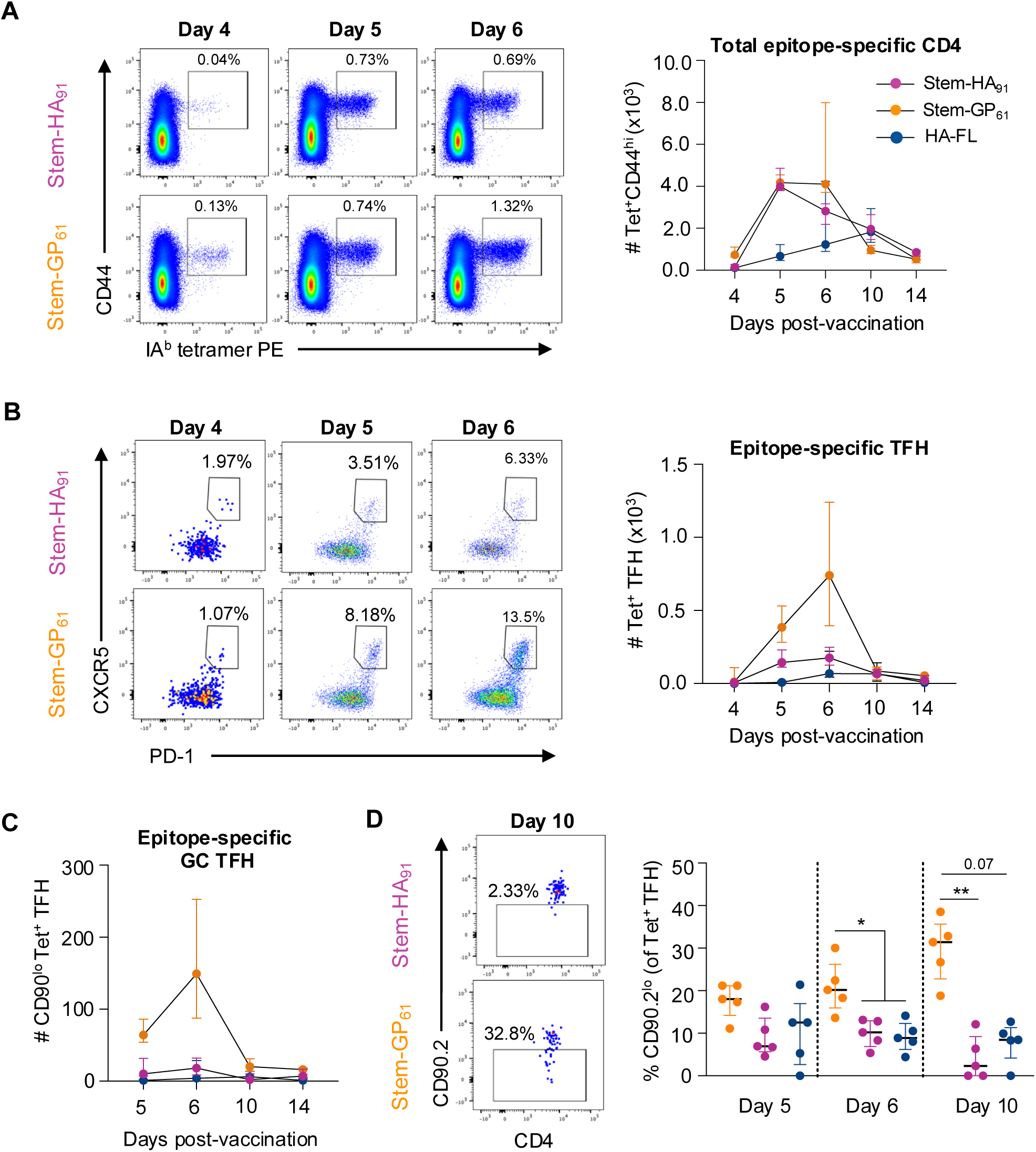
Differential GC recruitment of HA_91_ and GP_61_-specific TFH populations. **(A)** Longitudinal tracking of total, **(B)** TFH (CXCR5^hi^PD-1^hi^) or **(C)** GC resident TFH (CD90^lo^) antigen-specific CD4^+^ T cell numbers in the draining LN using IA^b^ tetramers. **(D)** Proportion of tetramer^+^ TFH with CD90^lo^ phenotype at days 5, 6 and 10 post-vaccination. Symbols indicate median and IQR (n=5 per group).

Recent work has shown that the identification of TFH based on high CXCR5 and PD-1 expression includes cells located both within and adjacent to the GC^26^. Downregulation of surface marker CD90 delineates a population of GC-resident TFH that emerge at day 5 post-immunisation and are dependent on MHCII-expressing B cells for their development and maintenance. Based on CD90 expression, we found significantly higher numbers of GC-resident TFH in stem-GP_61_ compared to stem-HA_91_ vaccinated animals from days 5-10 (Fig 4C). While this was partially driven by the overall greater number of GP_61_-specific TFH, the dynamics of CD90 downregulation also differed between HA_91_ and GP_61_ TFH populations. From day 5 to 10, the proportion of GP_61_ TFH with a GC-resident phenotype increased from a median of 18.0% to 31.4% (Fig 4D). In contrast, the proportion of HA_91_ TFH recruited into the GC declined from 6.9% on day 5 to 2.3% on day 10 (Fig 4D). As sustained GC B cell presentation of pMHCII is required to maintain CD90^lo^ TFH populations^26^, these data suggested that suboptimal class-II presentation of the HA_91_ peptide by B cells may limit the development of GC-resident TFH.

### Broader B cell presentation of HA_91_ supports the Stem-HA_91_ GC response

If differential B cell peptide-MHCII presentation controls the recruitment of ag-specific TFH into the GC, then altering B cell antigen presentation should rescue stem-HA_91_ immunogenicity (Fig 5A). Prior studies have suggested that defects in TFH differentiation can be overcome when DC antigen presentation is extended by boosting a protein immunization with cognate peptide three days later^27^. However, vaccination with 5ug of stem-HA_91_ supplemented with 5ug of free HA_91_ peptide at either day 0 or day 3 failed to rescue stem antibody titres (Fig 5B).

**Figure 5.**
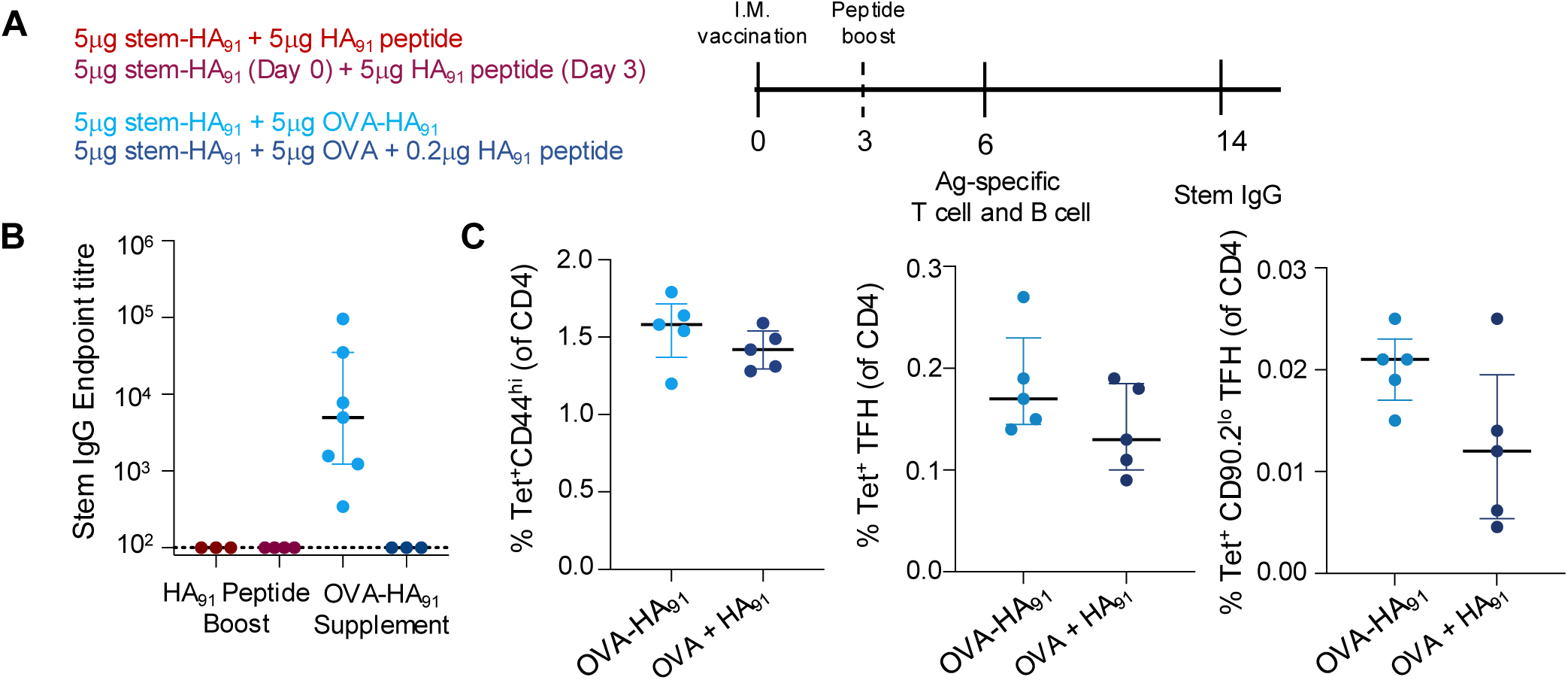
Modulation of stemHA_91_ immunogenicity through via altered B cell antigen presentation. **(A)** Overview of immunisation timeline and groups. (**B)** Stem IgG titres at day 14 post-vaccination for groups shown in panel A. N=3-4 per group from either one or two independent experiments. **(C)** Frequencies of total, TFH, or GC TFH HA_91_Tet+ T cells at day 6 post-vaccination with 5ug stem-HA_91_ + 5ug OVA-HA_91_ (light blue) or 5ug stem-HA_91_ + 5ug OVA + 0.2ug HA_91_ (dark blue). N=5 per group. Lines indicate median and IQR.

We next considered if the small pool of stem-specific naïve B cells may limit the presentation of pMHCII complexes, and contribute to the poor immunogenicity of stem-HA_91_ compared to HA-FL. We therefore immunised mice with 5ug of stem-HA_91_ co-formulated with 5ug of OVA-HA_91_ to increase the potential pool of naïve B cells presenting IA^b^/HA_91_. As controls for the OVA-induced GC response and increased dose of HA_91_ peptide, mice were vaccinated with 5ug of stem-HA_91_ co-formulated with 5ug OVA and an equimolar amount (0.2ug) of free HA_91_. Only animals vaccinated with stem-HA_91_ and OVA-HA_91_ exhibited stem-specific IgG at day 14 (Fig 5B), suggesting that covalent linkage of HA_91_ to an additional B cell antigen was necessary for T cell-B cell interactions that supported a productive stem-specific GC. Both groups exhibited similar HA_91_-specific T cell frequencies (Fig 5C). While frequencies of bulk HA_91_ TFH were slightly higher in the stem-HA_91_+OVA-HA_91_ group (median 0.17% of CD4 versus 0.11% for stem-HA_91_+OVA+HA_91_), CD90^lo^ GC resident TFH were 1.75-fold more frequent (0.021% versus 0.012%) (Fig 5C).

Collectively, these data suggest that the poor performance of the HA_91_ epitope as a sole source of CD4 T cell help is at least partially attributable to a failure to establish or maintain an antigen-specific CD90^lo^ GC-resident TFH population, which can be mitigated by expanding the pool of B cells capable of presenting IA^b^/HA_91_ complexes.

### Antigen dose differentially impacts CD4 priming and TFH differentiation

If differences in stem-HA_91_ and stem-GP_61_ immunogenicity were driven by pMHCII presentation in the GC, we next asked what the impact would be of reducing GP_61_ peptide availability. To titrate GP_61_ dose without changing the total amount of stem antigen available for B cell uptake, we vaccinated mice with varying doses of the stem-GP_61_ antigen supplemented with unmodified stem, such that the total dose of protein remained 5ug (Fig 6A). Draining lymph nodes were collected at day 6 post-vaccination, the timepoint of maximal GP_61_-specific T cell expansion and the appearance of stem-specific GC B cells. All immunization doses elicited a similar number of total GP_61_-specific CD44^hi^ CD4 T cells (Fig 6B), highlighting the marked efficiency of T cell priming by DCs.

**Figure 6.**
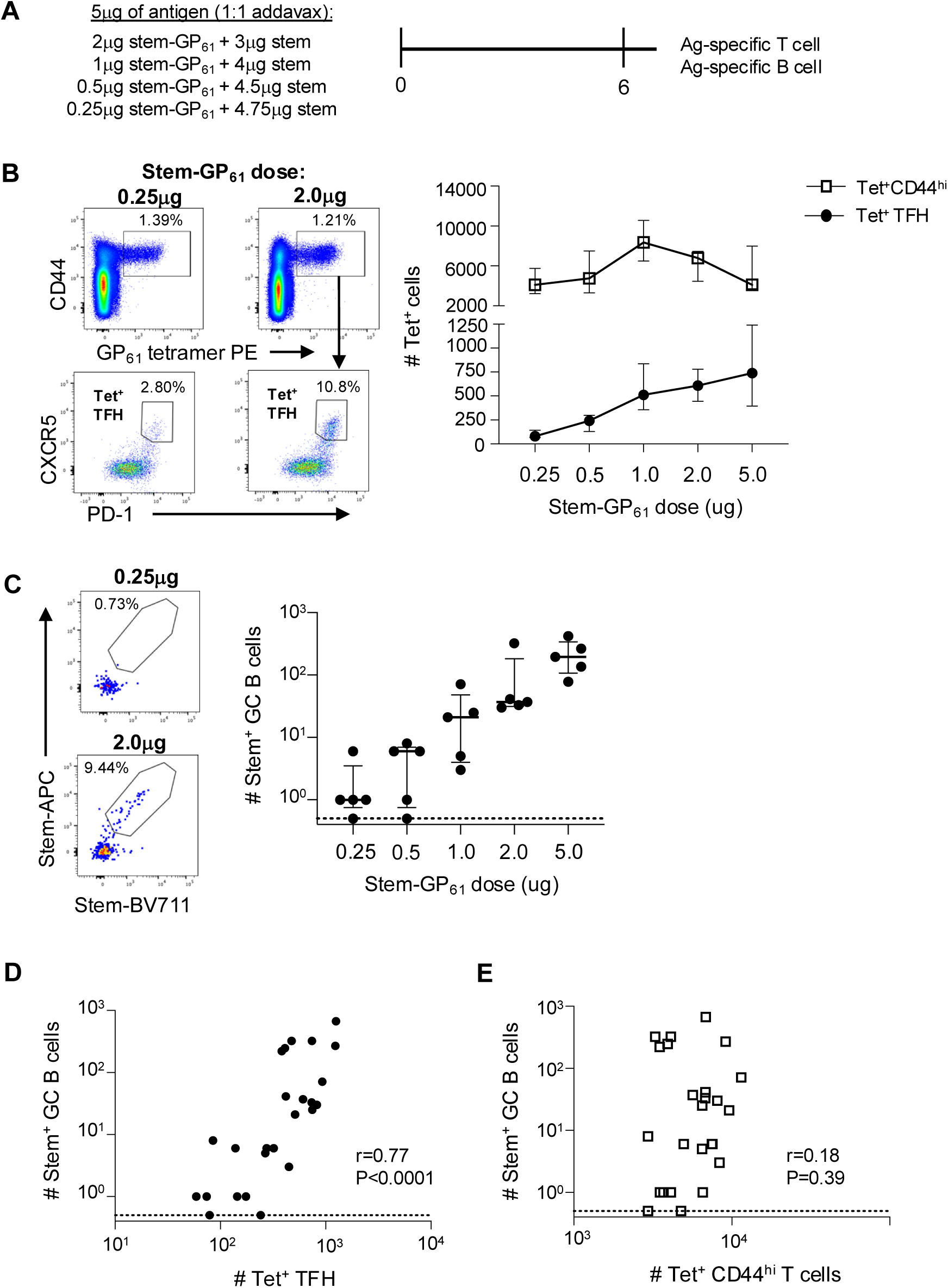
Modulation of stem immunogenicity through via altered B cell antigen presentation. **(A)** Antigen doses for titration of GP_61_ peptide availability. **(B)** Numbers of total and TFH GP_61_ Tet+ cells at day 6 post-vaccination (N=5 per group). **(C)** Number of stem-specific GC B cells at day 6 post-vaccination (N=5 per group). Symbols indicate median and IQR. **(D)** Correlation between number of Tet+ TFH or **(E)** total Tet^+^ cells with stem-specific GC B cells. Lymph nodes with no detectible stem-specific B GC B cells were arbitrarily assigned a value of 0.5 (marked with a dashed line) for graphical display. Statistics assessed by Spearman correlation.

In contrast, we observed a dose-dependent decrease in GP_61_-specific TFH differentiation (Fig 6B) that paralleled a dose-dependent decline of stem-specific GC B cells (Fig 6C). Accordingly, the number of stem+ GC B cells correlated with the number of ag-specific TFH but not total numbers of ag-specific CD4 T cells (Fig 6D-E), reinforcing that when T cell priming and TFH differentiation are decoupled, GC formation is controlled by TFH availability. Collectively, these data suggest that vaccine dosing sufficient to establish efficient B cell presentation of pMHC to pre-TFH cells is markedly higher than doses required for DC-priming of naïve T cells, and that differences in B cell presentation of HA_91_ and GP_61_ peptides may underpin the differential immunogenicity of stem-HA_91_ and stem-GP_61_ antigens.

## Discussion

Engineering vaccine immunogens to reduce inclusion of B cell epitopes with weak or narrow protective capability is an attractive approach for maximizing durable protection against challenging pathogens such as influenza, coronaviruses, HIV, and malaria. While such strategies facilitate highly tailored B cell engagement, increasing evidence suggests that a concomitant loss of CD4 T cell epitopes can hinder overall vaccine immunogenicity *in vivo*. Here, we assessed how CD4 T cell help supports productive GC reactions toward multimerized or soluble protein antigens, and found that T cell epitope specificity, rather than immunodominance, was a determining factor in TFH differentiation and vaccine immunogenicity.

Intrinsic protein immunogenicity is likely determined by the complex interplay of multiple factors including B cell precursor frequencies, antigen size, glycosylation, and the ability to elicit an effective CD4 T helper cell response. In a polyclonal system with soluble viral antigens, gross immunogenicity is likely to reflect the totality of available CD4 T cell help, with large antigens generally containing an increased number of CD4 epitopes than small antigens. This is consistent with the observed immunogenicity gradient of stem-ferritin nanoparticles across SMARTA, BL/6 and BALB/c mice; the broad ferritin-specific CD4 T cell pool in BALB/c mice was associated with 10-fold higher stem antibody titres compared to BL/6 animals. This appears to be generally consistent in genetically diverse human cohorts as well: large glycoproteins such as SARS-CoV-2 spike contain sufficient epitopes restricted by a broad array of HLA alleles to support immunogenicity in diverse populations. Small antigens like HA stem, SARS-CoV-2 RBD, HIV env or Plasmodium CSP, however, contain reduced numbers of CD4 cell epitopes that begin to restrict the quantity of help available in certain individuals or populations^15-18^. At this point, vaccine immunogens are at higher risk of failure, particularly for pathogens with high antigenic diversity^16^.

Considering these factors, it becomes essential to understand the minimal requirements for CD4 T cell help to support a robust GC reaction. Our data identify distinct scenarios under which vaccination can succeed or fail to generate a serological response. In the context of multimeric arrayed antigens, highly subdominant T cell help becomes sufficient to drive a vaccine-specific serological response. Despite using multiple assays, the frequency of stem- or ferritin-specific CD4 T cells elicited by primary vaccination of BL6 mice appears to be below our limit of detection. Nonetheless, the lack of stem-ferritin immunogenicity in SMARTA transgenic mice and ferritin-specific proliferative responses evident upon repeated boosting clearly support the contribution of low frequency or low affinity antigen-specific T cells to vaccine immunogenicity. Mechanistically, the augmented BCR crosslinking and NF-kB signaling induced by nanoparticle vaccines ^20^ may potentially reduce the magnitude and/or quality of CD4 T cell help required to initiate a stem-specific GC B cell response.

In the context of soluble non-arrayed antigens, we find that a single CD4 epitope is sufficient to support an antigen-specific GC reaction, but perhaps more importantly, find that only some epitopes drive CD4 T cell responses capable of providing suitable B cell help. The potent immunogenicity of the stem-GP_61_ protein clearly demonstrates how even a comparatively restricted CD4 T cell repertoire can support robust GC activity, as the naive repertoire of GP_61_-specific T cells is heavily biased toward TRAV14/TRAV14D and TRBV13/TRBV31 gene usage^28^. Our data therefore suggest that a highly diverse TFH pool may not be required for vaccine efficacy; rather, the rational selection of a small number of high-quality class II epitopes could be sufficient to support robust humoral immunity.

Collectively, our data support the development of GC TFH at an epitope-specific level by two distinct antigen presentation events: naïve T cell priming by DCs which drives immunodominance but not TFH selection, and subsequent T cell-B cell interactions that down select T cell specificities to populate the GC TFH pool. During the first 5 days post-vaccination, the kinetics of T cell expansion were comparable between HA_91_ and GP_61_ epitopes, likely reflecting naïve T cell precursor frequencies which are known to be high for GP_61_ compared to other epitopes such as OTII^28^. Interestingly, early DC priming was markedly efficient, as all doses of stem-GP61 antigen tested (0.25ug to 5ug) were saturating for T cell effector frequency, even when they compromised TFH numbers. It is currently unclear whether this is driven by differential antigen processing and presentation in B cells versus DCs, more efficient acquisition by DCs due to immunogen draining patterns in the LN, or a selective threshold imposed by low-frequency CD4 T cell/B cell interactions at the T:B border.

It is well known that the major selective event for TFH differentiation and GC residence is recognition of peptide:MHC complexes presented by B cells. However, our data suggest that even in a polyclonal context, there are stark differences in the capacity of epitope-specific T cell populations to provide B cell help. Mechanistically, this appears to be governed by a low propensity of some polyclonal populations (like HA_91_-reactive T cells) to undergo differentiation into CD90lo GC-resident TFH. Our panel of stem-peptide immunogens identified some ‘immunocapable’ epitopes (GP_61_ and HA_523_), which support GC responses using the endogenous T cell and B cell repertoire in BL/6 mice. Other epitopes, such as HA_91_, required manipulation of B:T interactions (e.g. expansion of the peptide-presenting B cell pool) to effectively drive GC. Possible mechanisms underpinning these observations include peptide-intrinsic differences in presentation/processing by B cells (but not DCs) or epitope-specific differences in TCR/MHCII avidity at an aggregate, polyclonal level. The linear relationship between dose of stem-GP_61_ and GP_61_ TFH differentiation suggest the absolute densities of IAb/GP_61_ on the surface of stem-specific B cells dictates a propensity to recruit GP_61_ T cells into the GC, potentially favouring a deterministic role for B cell peptide presentation in controlling TFH composition.

Additional study is required to extend these observations into human cohorts. Despite the incredible diversity provided by HLA polymorphism and TCR repertoires, mounting evidence suggests that the human TFH repertoire may also be relatively restricted, with individual epitopes or class II alleles exerting substantial impacts on the outcome of vaccination^22,29^. Antigens such as HIV env, *Plasmodium* CSP and SARS-CoV-2 RBD frequently contain only 1-2 epitopes that are recognized on an individual level^15-18^. The qualitative differences between HA_91_ and GP_61_ highlight the potential pitfalls of relying on few, endogenous epitopes for consistent vaccine immunogenicity. Further work is required to clarify whether multiple, subdominant epitopes provide additive or synergistic support for GC B cell development. These data also underscore the need to accurately predict the quality of B cell help, rather than just immunogenicity or immunodominance, of any given epitope and responding T cell population to aid vaccine design efforts.

## Methods

### Mouse immunizations and infection

C57BL/6, BALB/c, and SMARTA-transgenic (C57BL/6 background) mice were bred in-house under specific pathogen-free conditions in the animal facility at the Peter Doherty Institute of Infection and Immunity, University of Melbourne, Australia. Mouse studies were carried out in accordance with the University of Melbourne Animal Ethics Committee (no. 22954). All mice were female and aged 6 to 12 weeks at the time experiments commenced. A total of 5μg (unless otherwise indicated) of protein, peptide, or nanoparticle were formulated in phosphate buffered saline (PBS) at a 1:1 ratio with Addavax (InvivoGen, cat#INV-vac-adx-10) adjuvant in a total volume of 100μL. Mice were anesthetized by isoflurane inhalation with oxygen flow at 2L/min and isoflurane vaporizer set to 3 (Stinger Anesthetic Machine), prior to 50μl intramuscular injections at the left and right quadriceps using a 29G needle. For influenza infections, mice were anesthetized as above and intranasally infected in a volume of 50μl with a sublethal dose of 50 TCID_50_ of A/Puerto Rico/8/34 (PR8). At experimental endpoints, mice were killed using CO_2_ asphyxiation using a delivery of 50% chamber volume per minute.

### Protein expression

Stem and Stem-Ferritin immunogens were prepared in-house, as previously described^9,30^. Briefly, stabilized HA stem proteins were engineered for A/Puerto Rico/08/1934 using methods established previously for the design of Gen6 HA stem in Yassine et al.^24^. Stem-Ferritin nanoparticles were expressed by transient transfection of Expi293F (Life Technologies, Thermo Fisher Scientific) suspension cultures and purified using ion exchange chromatography with HiTrap Q HP column (GE Healthcare) and exclusion chromatography.

For stem proteins conjugated to CD4 T cell epitopes, expression constructs substituting the original Avitag for peptide sequences for OTII (ISQAVHAAHAEINEAG), HA_91_ (RSWSYIVETPNSENGIC), HA_115_ (YEELREQLSSVSSFERF), HA_301_ (AINSSLPYQNIHPVTIG), HA_523_ (SMGIYQILAIYSTVASS) and GP_61_ (GLKGPDIYKGVYQFKSVEFD) were synthesized (GeneArt), cloned into mammalian expression vectors and expressed via transient transfection of Expi293 suspension cultures (Life Technologies, Thermo Fisher Scientific). Proteins were purified by polyhistadine-tag affinity chromatography and gel filtration.

### Enzyme-linked immunosorbent assay (ELISA)

Blood samples for serum isolation were collected either by submandibular bleed or terminal cardiac puncture bleed using a 26G needle. 96-well MaxiSorp plates (ThermoFisher, cat# 3442404) were coated with 2μg/mL recombinant stem protein overnight at 4°C. Plates were washed with 0.05% (v/v) Tween20 (Sigma, cat# P1379) + PBS and blocked with 1% (v/v) FCS in PBS for 1 hr, room temperature, before incubation with serial dilutions of sera for 2 hr. Plates were washed and horseradish peroxidase-conjugated anti-mouse IgG (1:15 000; Seracare, cat# 5450-0011) was added for 1 hr. After washing, plates were developed with 3,3′,5,5′-Tetramethylbenzidine (TMB; ThermoFisher, cat#SB02), stopped with 0.16M sulfuric acid (Sigma, cat# 84727) and the absorbance measured at 450nm on FLUOstar Omega microplate reader (BMG Labtech). Curves were fitted (four-parameter log regression) and end-point titrations calculated as the reciprocal serum dilution yielding 2x background using GraphPad Prism version 10.

### Generation of B cell probes and peptide MHC II tetramers

Recombinant biotinylated stem protein biotinylated using BirA (Avidity) and stored at –80°C. Conjugation was performed by sequential addition of streptavidin-PE or -APC (Life Technologies, cat#S866 and S868) or streptavidin-BV711 (BD, cat# 563262). Mouse H2-IAb RSWSYIVETPNSENGI PE-conjugated tetramer (IAb/HA_91_) was generated by ProImmune. Biotinylated mouse H2-IAb DIYKGVYQFKSV monomer (IAb/GP_61_; ProImmune) was tetramerized by sequential addition of streptavidin-PE (Life Technologies, cat# S866).

### Detection of antigen-specific B and T cells ex vivo

Iliac and inguinal vaccine-draining lymph nodes were passed through a 70μm filter, centrifuged (500*g*, 7 min) and washed with PBS prior to viability staining with live/dead red for 3 min (Invitrogen, cat# L34972). Cells were Fc blocked with anti-CD16/32 antibody (BioLegend, cat# 101302) for 10 min and stained with surface antibodies of interest for 30 min at 4°C. Cells were then washed and fixed with 1% (v/v) formaldehyde (BD, cat# 554655) before acquisition on a BD Symphony or Fortessa flow cytometer. For detection of antigen-specific B cells, PE, APC or BV711 conjugated stem probes were included in the surface staining antibody cocktail. For detection of antigen-specific T cells, single-cell suspensions were pre-incubated at 37°C with 50nM dasatinib (Sigma, cat# SML2589) in 2% (v/v) FCS in PBS containing 1μg/mL anti-TCRý (BD, cat# 553167). After 30 min, 4μg/mL of IAb/HA_91_ or IAb/GP_61_ tetramers were added for a further 3 hr, before proceeding with viability and surface staining. Data were analyzed using FlowJo (v10.10, BD Biosciences). Data collection and analysis were not performed blind to the conditions of the experiments.

### Activation Induced Marker assays

Individual HA, stem (BEI Resources) or ferritin (GenScript) peptides (15-mer with 11 amino acid overlap) were pooled and used to detect antigen-specific CD4+ T cell responses in vitro. LN single cell suspensions were cultured in 96-well round-bottom plates in RPMI-1640 (Thermo Fisher Scientific) with 10% (v/v) FCS and 2% (v/v) penicillin-streptomycin (RF10) media containing 20mM A438079 (Santa Cruz, cat# 203788), anti-mouse CD154 antibody and peptide pool of interest (2mg/mL/peptide) or an equivalent volume of DMSO (Sigma, cat# D1435). Following 18hr of stimulation, cells were washed in PBS, stained for viability (3 minutes at room temperature) and stained with surface antibodies of interest.

### T cell proliferation assay

To generate single-cell suspensions, spleens were passed through a 70μm filter and centrifuged (500xg, 7 min). Red blood cells were lysed using 1x BD PharmLyse (Cat #555899) for 3 min, and the reaction quenched with 1x PBS. Splenic single-cell suspensions were labelled with 2.5mM of CellTrace Violet (CTV; Invitrogen, cat#C34557) for 10 min and washed twice (500xg, 7 min) with RF10. 4×10^5^ CTV-labelled cells were cultured in 96-well round-bottom plates in RF10 with 0.5mg/mL HA, stem or ferritin peptide pools, or anti-CD3/CD28 Dynabeads (Gibco, cat # 11456D) for 4 days at 37°C, prior to staining for flow cytometric analysis.

### Immunofluorescent microscopy

Fresh tissues were snap-frozen in Tissue-Tek O.C.T. compound (Sakura Finetek USA) and stored at -80°C. 7 um sections were cut using the Leica CM3050S cryostat (Leica Biosystems). Prior to staining, sectioned tissues were fixed in cold acetone solution (Sigma) for 10mins, rehydrated with PBS for 10mins, and then blocked in 5% (w/v) bovine serum albumin (Sigma) and 2% (v/v) normal goat serum (Jackson ImmunoResearch). A cocktail of antibodies including GL7 AF488 (clone GL7, BioLegend) and B220 BV421 (clone RA3-6B2, BioLegend) were added for 1 hr at RT. Slides were mounted with ProLong Diamond Antifade Mountant (Life Technologies). Tiled z-stack images at 20x magnification and 1 airy unit were acquired on a LSM780 microscope (ZEISS) and analyzed with Fiji software^31^.

### Statistical analysis

Data are presented as median ± interquartile range. All statistical analysis was performed in Prism 10 (GraphPad) using nonparametric statistical tests as indicated (making no assumptions about data normality). P<0.05 was considered statistically significant.

## Acknowledgements

This work was funded by an NHMRC Ideas grant to JAJ. HXT, AKW and JAJ are supported by NHMRC Investigator Grants, and JAJ is supported by the Sylvia and Charles Viertel Senior Medical Research Fellowship. The funders had no role in study design, data collection and analysis, decision to publish or preparation of the manuscript.

## Supplementary Figures

**Supplementary Figure 1.**
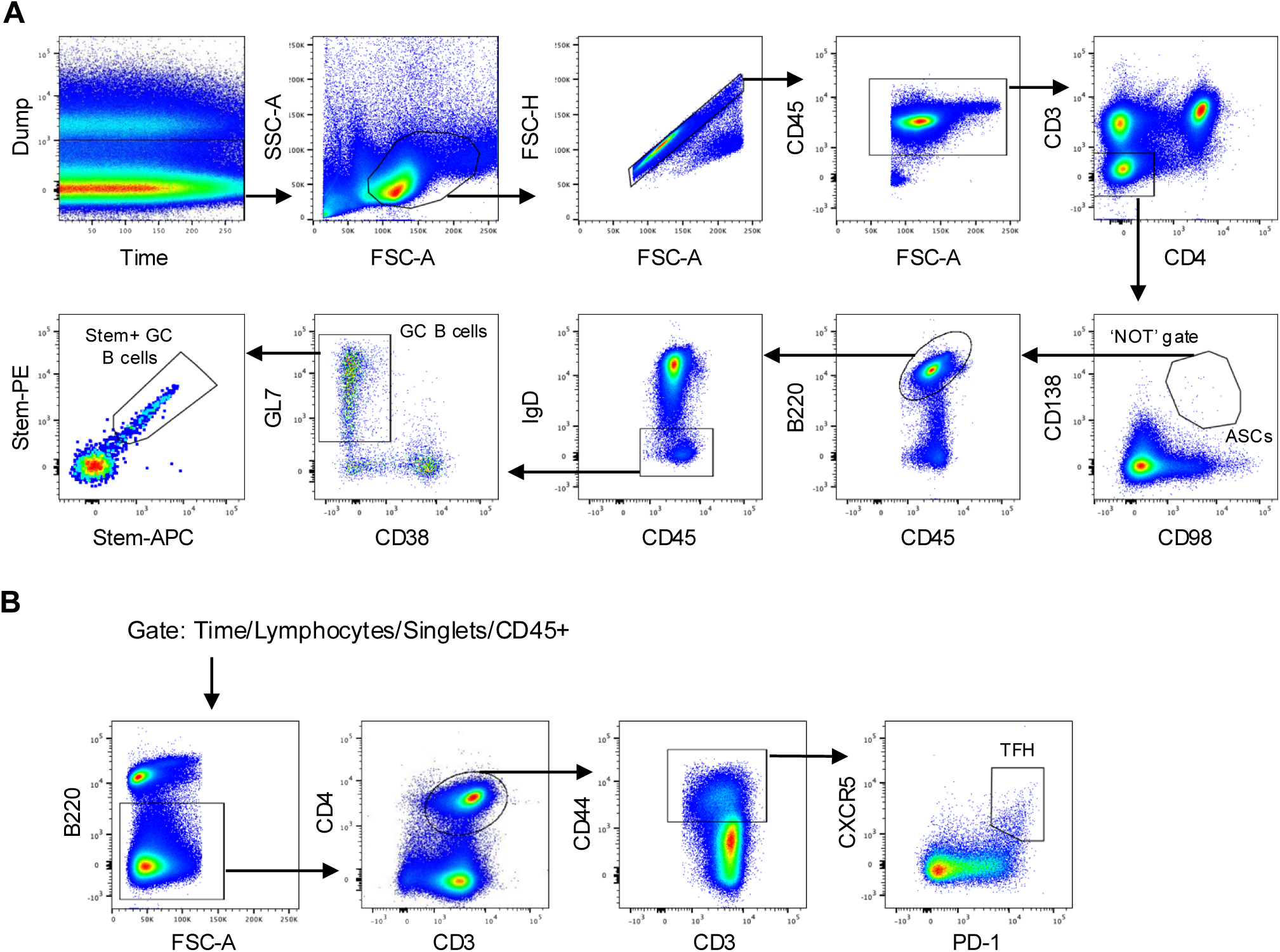
Gating strategy for GC B cell and TFH populations. **(A)** Lymphocyte populations are gated through a Dump (F4/80 and live/dead) vs Time plot, then identified by forward and side scatter, with exclusion of doublets (FSC-A vs FSC-H) and selection of CD45+ cells. Antibody secreting cells are defined as CD3 -CD4-cells that co-express CD138 and CD98. Class-switched B cells are defined as non-ASCs that are B220+ and IgD-. Germinal centre B cells are gated as GL7+CD38lo, and antigen specificity determined by staining with recombinant protein tetramers. **(B)** CD4+ T cells are identified as B220- and CD3+CD4+. Antigen-experienced cells are gated as CD44hi, and TFH are defined as CXCR5+PD-1+ cells.

**Supplementary Figure 2.**
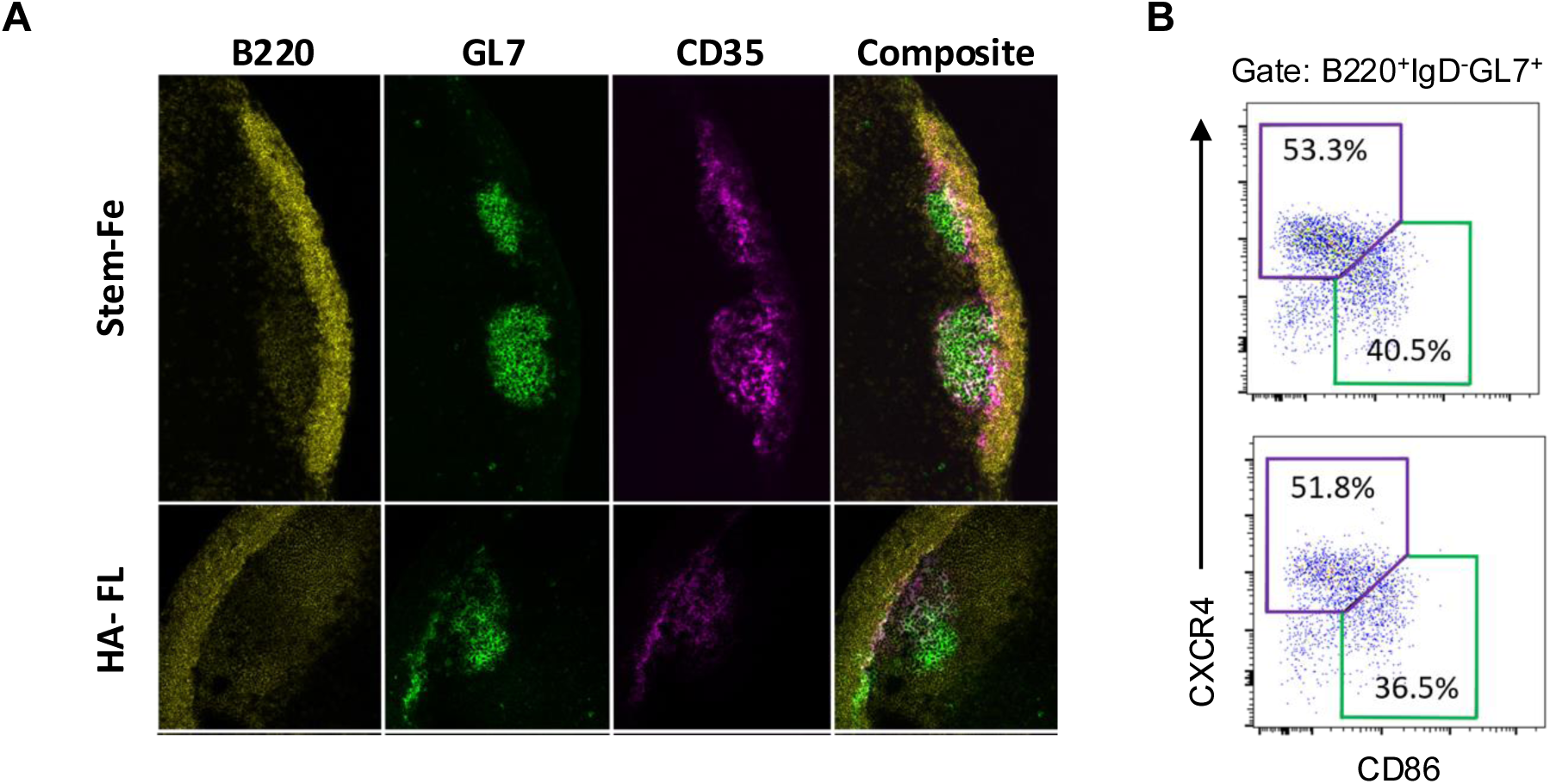
Germinal centre morphology following stem-Fe vaccination. (A) Confocal imaging of vaccine draining lymph nodes showing localisation of B220^+^ B cells (yellow), GL7^+^ GC B cells (green), and CD35+ follicular dendritic cells (magenta). (B) Identification of light zone (CXCR4^lo^CD86^hi^) and dark zone (CXCR4^hi^CD86^lo^) GC B cells in stem-Fe (top) or HA-FL (bottom) vaccinated animals. Data are representative of 5 individual animals in each group.

**Supplementary Figure 3.**
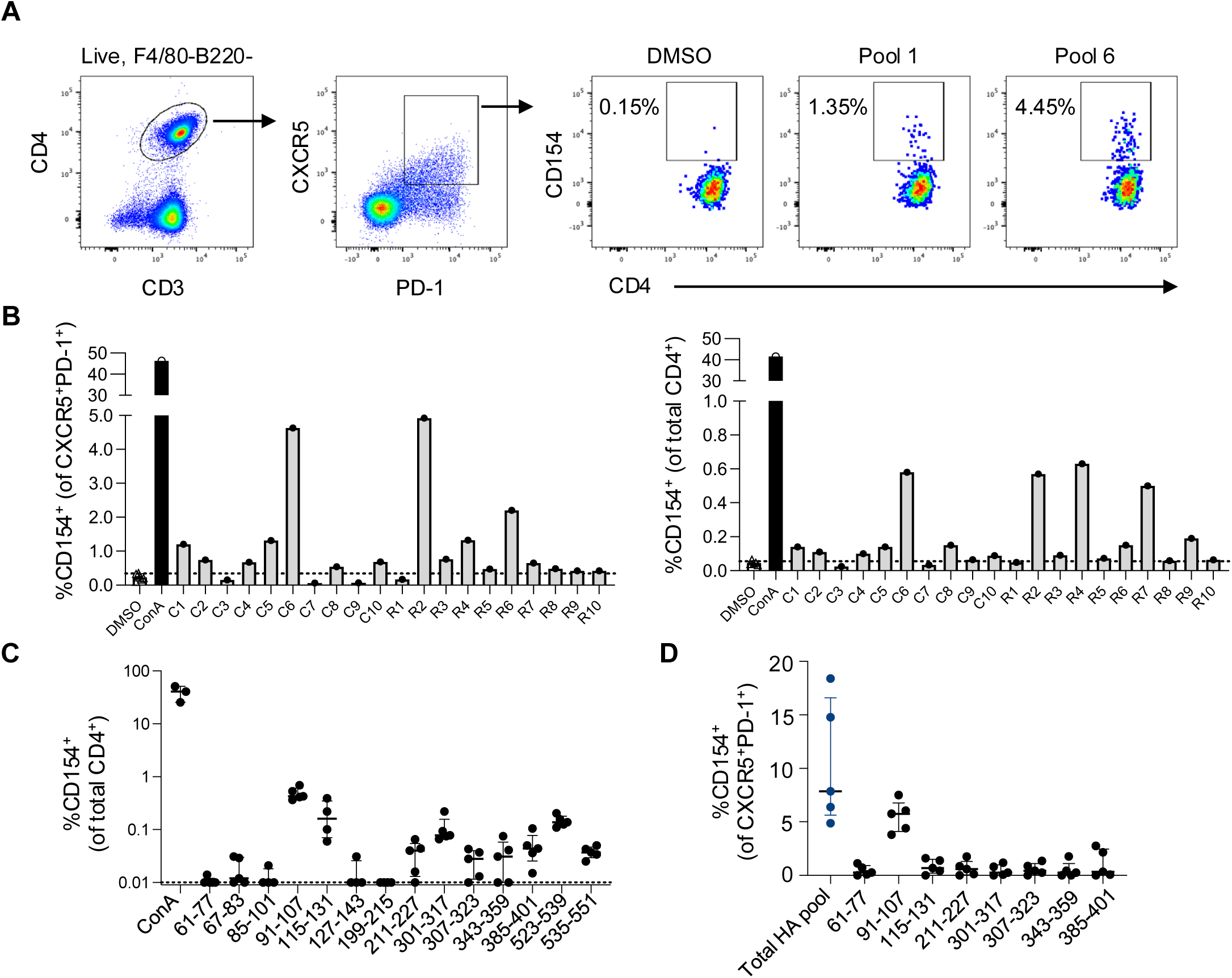
Identification of immunogenic peptides in PR8 HA. **(A)** Representative staining of CD154 upregulation on CXCR5^+^PD-1^+^ CD4^+^ T cells in response to *in vitro* stimulation with peptide pools covering the PR8 HA protein. **(B)** Frequencies of peptide-specific or ConA-responsive cells among CXCR5^+^ (left) or total (right) CD4 T cells following PR8 infection. **(C)** Frequencies of peptide-specific bulk CD4^+^ T cell responses in the mediastinal LN at day 14 post-PR8 infection. Each dot represents a single mouse (n=4-5 per group). **(D)** Peptide- or pool-specific TFH responses in draining LN following intramuscular vaccination with soluble PR8 HA antigen.

**Supplementary Figure 4.**
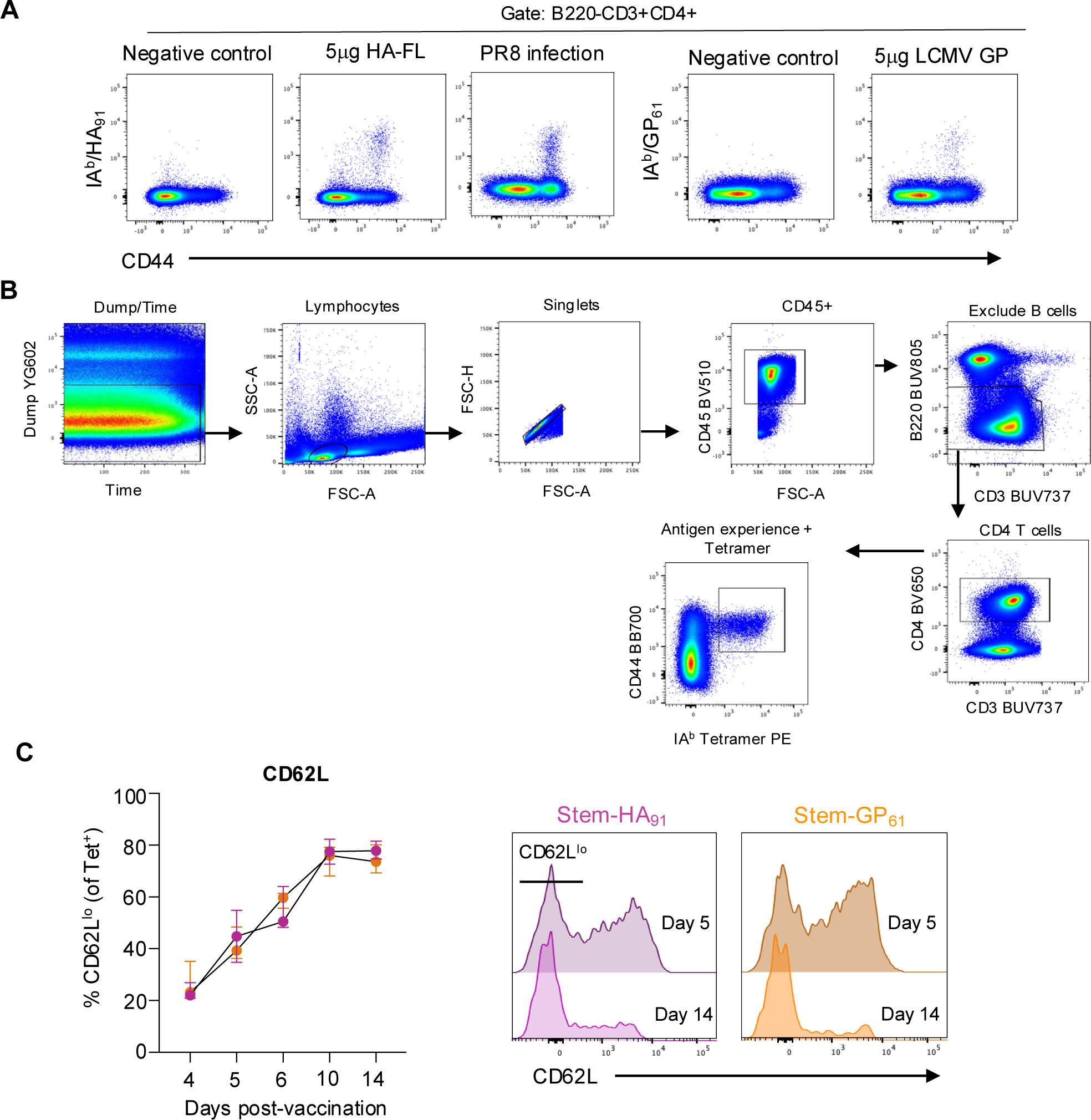
Gating strategy for longitudinal tracking of antigen-specific CD4^+^ T cells. **(A)** Representative staining of IA^b^/HA_91_ and IA^b^/GP_61_ tetramers in draining lymph nodes at day 14 following HA-FL vaccination, PR8 infection or LCMV GP vaccination. **(B)** Antigen-specific cells were defined as Live, F4/80-CD45+B220-CD3+CD4+CD44hi tetramer+ lymphocytes. **(C)** Expression of CD62L on Tet^+^ cells from days 4-14 post-vaccination. Graph indicates median and IQR, N=5 per group.

